# Pathogenesis of H5N1 Clade 2.3.4.4b in dry Jersey cows following intramammary inoculation shows within-host compartmentalization

**DOI:** 10.64898/2026.03.04.709389

**Authors:** Konner Cool, Jessie D. Trujillo, Taeyong Kwon, Gagandeep Singh, Sujan Kafle, Chester D. McDowell, Isaac Fitz, Shanmugasundaram Elango, Eulim Lyoo, Govindsamy Vediyappan, Wanting Wei, Heather M. Machkovech, Franco Matias Ferreyra, William C. Wilson, Brittany Cronk, Igor Morozov, Thomas C. Friedrich, Diego G. Diel, Natasha N. Gaudreault, Juergen A. Richt

## Abstract

Dairy cattle have emerged as a prolific amplifying host for highly pathogenic avian influenza virus (HPAIV) H5N1 clade 2.3.4.4b and a new source for cross-species and zoonotic transmission. Independent introductions of H5N1 with unclear exposure routes have been reported in several dairy herds across the U.S. These events escalate the pandemic potential of HPAIV H5N1 as transmission within and between mammalian species present opportunities for mammalian adapted H5N1 viruses to emerge. Although more than 1000 herds have been infected, bovine H5N1 influenza virus pathogenesis, transmission, and evolution in dairy cattle remains not well characterized. Working with H5N1-infected lactating cattle in high containment has been a major challenge due to the required infrastructure and logistics associated with housing, husbandry, and waste management for this model. Thus, developing alternative bovine models that maintain biological relevance while reducing operational complexity is warranted. Here we evaluate the susceptibility of lactating Jersey cattle in the dry-off period and characterize the effect of inoculation dose on the mammary pathogenicity of HPAIV H5N1 genotype B3.13. The results of this study demonstrate that dairy cows 21 days into the dry-off period are highly susceptible to HPAIV H5N1, recapitulating the severe clinical and pathological outcomes observed in infected lactating cows under experimental conditions and in field cases. We also observed an association between virus dose and the onset and severity of mastitis in individual udder-quarters. Overall, this study demonstrates that dry cows can provide a feasible model to study H5N1 virology, pathology, and humoral immunology in dairy cows.

## Introduction

The global panzootic caused by highly pathogenic avian influenza virus (HPAIV) H5N1 clade 2.3.4.4b has affected hundreds of species across five continents, with fatal outcomes reported in wild carnivores and marine mammals ^1–3^. Introduced to North America in late 2021, viruses carrying hemagglutinin (HA) of clade 2.3.4.4b have become enzootic in wild and aquatic bird species with frequent spillover into commercial poultry. Distinct from previous HPAIV outbreaks, the host range of clade 2.3.4.4b viruses has expanded to include numerous mammalian and peridomestic species, escalating concerns for the emergence of mammalian adapted H5N1 strains with pandemic potential.

Outbreaks of HPAIV H5N1 in lactating dairy cattle were first reported in March of 2024, expanding to 19 U.S. states by the end of 2025. Infected dairy herds have become a primary source for human exposure to H5N1 in North America as the number of confirmed zoonotic transmission events associated with exposure to infected dairy cows is nearly twice that of exposure to infected poultry^4,5^. H5N1 infected dairy herds have also been associated with infection of cats, poultry flocks, and other peridomestic species in proximity to infected dairy parlors, possibly through exposure to raw milk and fomites ^6–9^. Spillover into dairy cows is believed to be direct from wild avian, although the circumstances surrounding initial exposure/infection is not fully understood yet. At least three additional independent introductions of HPAIV H5N1 (genotype D1.1) have been reported in dairy herds located in Nevada, Arizona, and Wisconsin in 2025^10^. This shift in host range of influenza has made domestic cattle a novel and understudied host, posing new challenges for biosecurity, animal health, and *One Health* preparedness.

Compared to other mammalian hosts, infection and pathogenesis of H5N1 in dairy cattle is atypical by manifesting itself primarily in the udder (mammary tissue), with limited tropism to other tissues. Clinical signs in infected dairy cattle usually present as acute with significant reduction in milk production and milk quality, reduced rumination (inappetence), pyrexia, and mastitis. Virus replication is prolific in the bovine udder and shedding occurs primarily through milk, often reaching up to 10^9^ TCID_50_/mL in milk. Transmission between cows seems efficient in most dairy operations, likely occurring during the milking process, or through contaminated equipment and farm practices. Alternative routes of intra-herd transmission have been suggested to occur via *i)* aerosolized virus in milking parlors, *ii)* directly from cow-to-cow (e.g., milk snatching), *iii)* indirectly through fomites, run-off, contaminated water and feed, care-takers, peridomestic species, and wind or *iv)* mechanical transmission via house flies (*Musca domestica*), blow flies (*Calliphoridae*), or wildlife (rodents, swallows); these alternative routes cannot be ruled-out as they remain to be fully elucidated ^11–17^. The widespread dissemination of HPAIV H5N1 between dairy herds and States has been linked primarily to transportation of infected cattle, contaminated equipment, and/or vehicles, although alternative transmission chains have been proposed as described above.

Experimental infection models have characterized HPAIV H5N1 clade 2.3.4.4b infection in lactating cattle ^11,18–21^. These initial studies successfully recapitulated the clinical and pathological disease observed in natural infections. However, significant gaps remain in understanding the susceptibility, pathogenesis, and evolution of H5N1 in this host. Additional studies are needed to address these gaps, but there is a limited capacity in presently available animal biosafety level-3 agricultural (ABSL-3Ag) containment facilities to conduct *in vivo* experiments with HPAIV H5N1 in lactating cattle. Primary constraints to this type of research are in the lack of necessary infrastructure to accommodate and adequately care for such a large species, including management and disposal for the amount and type of waste that is produced.

The aim of this study was to evaluate dairy cows in the dry-off period as an alternative model to examine H5N1 pathogenesis, virus replication, shedding, and immunogenicity following intramammary infection. Drying off dairy cows involves stopping the milk production cycle roughly 45 to 65 days before the next calving. This provides a necessary rest period for the cow to repair mammary tissue, improve health, and boost future milk production. Drying off is achieved by either abruptly ceasing milking or gradually reducing milking frequency, often supported by lower-energy feed, teat sealant application, and antimicrobial therapy. While cows at this stage in production can produce low levels of milk, they do not require daily milking. By enlisting cows in the dry-off period, several constraints posed by the required ABSL-3Ag infrastructure and husbandry associated with milking are reduced. We demonstrate here that clinical and pathological disease, virus replication, and host response in dry cows is similar to that of lactating cows following intramammary inoculation. Furthermore, we describe the compartmentalized, dose-dependent pathogenicity observed during H5N1 virus infection in the udder of dry cows.

## Results

### HPAIV H5N1 clade 2.3.4.4b causes severe clinical disease in dry cows

Six monoparous cows (Jersey breed) entering the dry off period were used in this study. All cows were in good physical condition, seronegative for influenza A-specific antibodies, and PCR-negative for most of the common bovine respiratory disease complex (BRDC)-associated pathogens (see **Supplementary Table 1**). After a 14-day acclimation period, cows were monitored to establish relative baselines for feed intake, rectal temperature, and various other clinical criteria (see **Supplementary Table 6**). To evaluate the effect of virus dose on the clinical course and severity of disease in individual udder quarters, each cow was inoculated intramammary with 0.5 mL of a virus suspension per quarter containing either 10^3^ (left hind quarter), 10^4^ (right front quarter), or 10^5^ (right hind quarter) TCID_50_ of HPAIV H5N1 (genotype B3.13); one quarter (left front) was mock inoculated with cell culture media and served as a negative, mock control (see **Supplementary Figure 1**).

#### Clinical manifestation and early convalescence

Clinical signs were observed in all (5/5) inoculated cattle within 48 hours post-inoculation. Acute clinical signs included transient marked reduction in feed intake (inappetence) and pyrexia. Rectal temperatures exceeding 105°F (normal temperature for cows is 100.4°F) were recorded in all (5/5) cows at 2 DPI and remained elevated (>103°F) in 3/5 cows at 3 DPI (**Figure 1**). Although randomization was initially used to designate cows for serial euthanasia at pre-determined end-points, selection for euthanasia was ultimately determined by clinical criteria and humane endpoints. The cow euthanized at 3 DPI (cow #89) was selected based on the relative rate and degree of clinical deterioration compared to the other cows; cow #89 exhibited labored breathing, warm and swollen udder quarters, dark tacky stool, and lethargy. Similar clinical signs were presented in the cow euthanized at 4 DPI (cow #93), with the addition of blood-tinged vaginal discharge, a presumptive indicator of post estrus status. Between 4-6 DPI, rectal temperatures and appetite suggested that cows were recovering from the acute viral infection. However, clinical signs returned by 7 DPI, entering into a second stage of inappetence and pyrexia with further swelling of the udder and teats, accompanied by tenderness, meeting the criteria for humane endpoints at 7 and 10 DPI.

**Figure 1:**
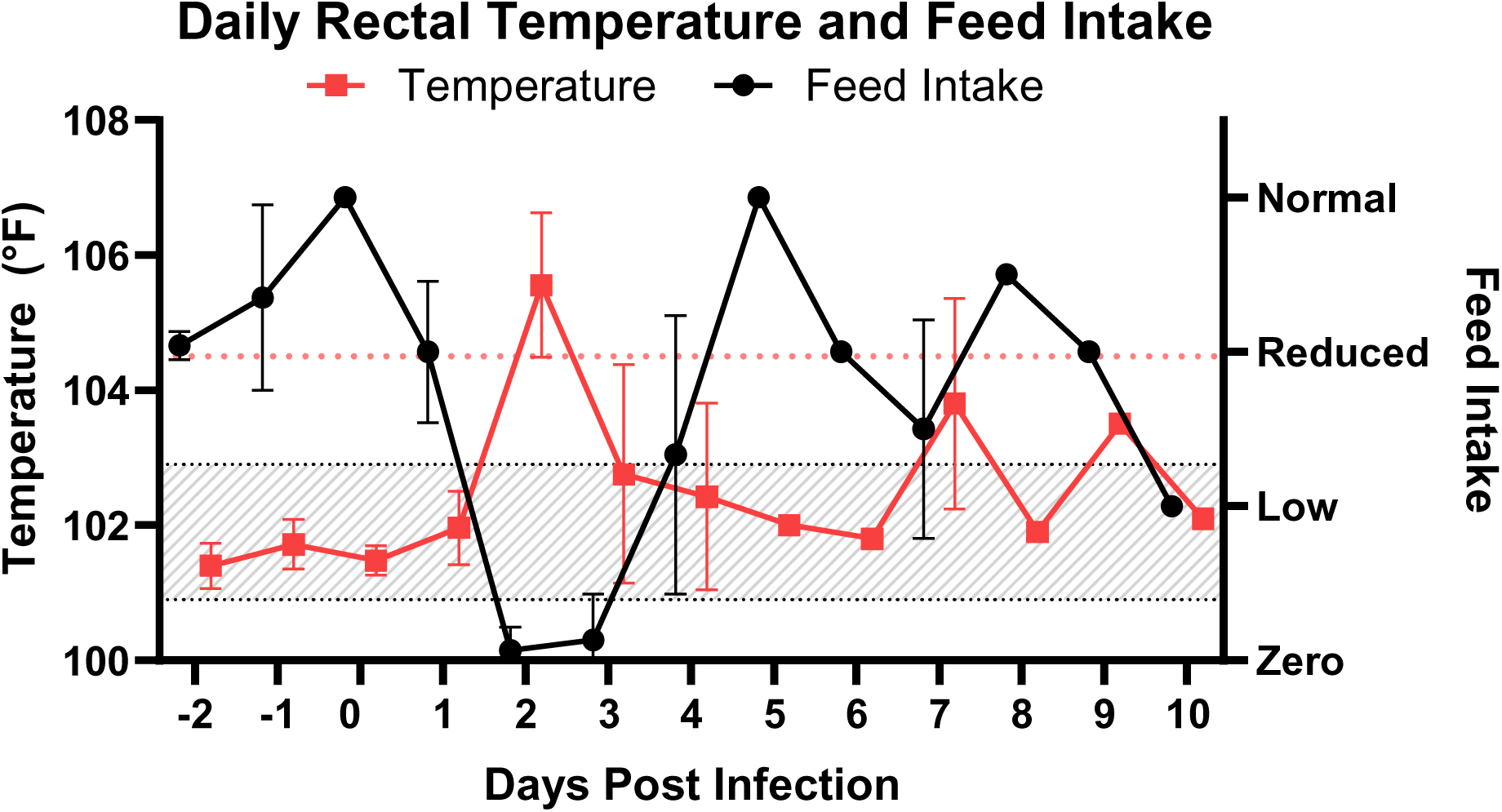
Daily rectal temperature and feed intake. Rectal temperature (°F), plotted on the left Y-axis, are presented as the mean (red square) and standard deviation (red vertical bars) of all recorded temperatures from each day (X-axis) throughout the study period. The gray shaded area represents the expected normal temperature (homeostatic) range for adult Jersey cows, whereas the red dotted line is used to indicate high fever. Plotted on the right Y-axis is a qualitative measure of observed feed intake. Each data point (black circle) and error bar (black vertical line) represents the mean and standard deviation, respectively, of feed intake for all cows at specified time-points.

#### Dose-dependent progression of viral mastitis

Clinical observations and standardized scoring with the California Mastitis Test (CMT) were recorded daily to document physical appearance of individual udder quarters and changes in milk quality. The onset and severity of mastitis was H5N1 virus dose-dependent and remained compartmentalized by quarter. Physical changes in individual quarters and milk quality aligned with CMT scores and virus loads shed in milk for the corresponding udder quarters. These changes were consistent between different cows and udder quarters inoculated with the respective H5N1 virus doses.

Mock inoculated quarters displayed little or no overt clinical signs of mastitis, and CMT scores remained low (within normal limits) until 7 and 10 DPI. Virus inoculated quarters were clinically distinct from one another, most apparent between low-dose (10^3^ TCID_50_) inoculated and high-dose (10^5^ TCID_50_) inoculated quarters (**Supplementary Figure 3**). Udder quarters that were inoculated with mid (10^4^ TCID_50_) - or high-doses of the virus displayed early overt clinical signs of mastitis, including erythema, swelling, and heat, and produced discolored, clotted milk as early as 2 DPI (cow #89, 90, 22) (**Supplementary Figure 4**). These quarters also exhibited the most rapid escalation in CMT scores (**Supplementary Figure 2**). Less severe symptoms and delayed CMT positivity were observed in the low-dose–inoculated quarters.

### Virus shedding in milk/mucosal surfaces and antibody response

Virus was recovered from milk in the low-, mid-, and high-dose inoculated quarters from 1-7 DPI, reaching peak titers of 10^8-^10^9^ TCID□□/mL at 2 DPI (**Figure 2**). Virus loads in milk at 1 DPI corresponded to the virus dose administered to respective quarters but converged across quarters by 3 DPI. Virus was recovered from the milk of all inoculated quarters for each cow remaining until 8 DPI. Throughout the entire study period, virus titers in milk from low-dose quarters inoculated quarters remained generally lower compared to the higher dosed quarters. Markedly lower titers of virus were also recovered from the milk collected from uninoculated quarters of two cows at 2 DPI, and a third cow at 3 DPI, but not on any day thereafter. As outliers, these data could suggest that samples were most likely contaminated during collection due to the high (peak) levels of virus shedding occurring at this time from the other three inoculated quarters. In addition, oral, nasal, rectal, and external teat swabs were collected daily to test for virus shedding. Swabs of the vaginal mucosa were collected daily starting at 4 DPI upon the observation of vaginal discharge containing blood in cow #93. Influenza A virus (IAV) RNA was detected at one or more collection time-points in all mucosal sample types. IAV-RNA was most consistently detected in external teat swabs, followed by vaginal swabs, oral swabs, and nasal swabs (**Supplementary Figure 5; Supplementary Table 5**). The highest loads of IAV-RNA (low Ct) were present in teat swabs throughout the entire study period, with IAV-RNA shedding frequently detected in nasal and oral swabs collected at 2 DPI (**Supplementary Table 5**). Infectious virus was recovered primarily from teat swabs, and from a single oral swab at 2 DPI (cow #22); all other mucosal swabs yielded RNA below infectious thresholds. No IAV-RNA was detected in whole blood samples during the study period. The data so far indicate robust virus replication in the udder; however, spread to other tissues seems to have occurred, but generally virus is present at very low levels near the threshold of vRNA detection.

**Figure 2:**
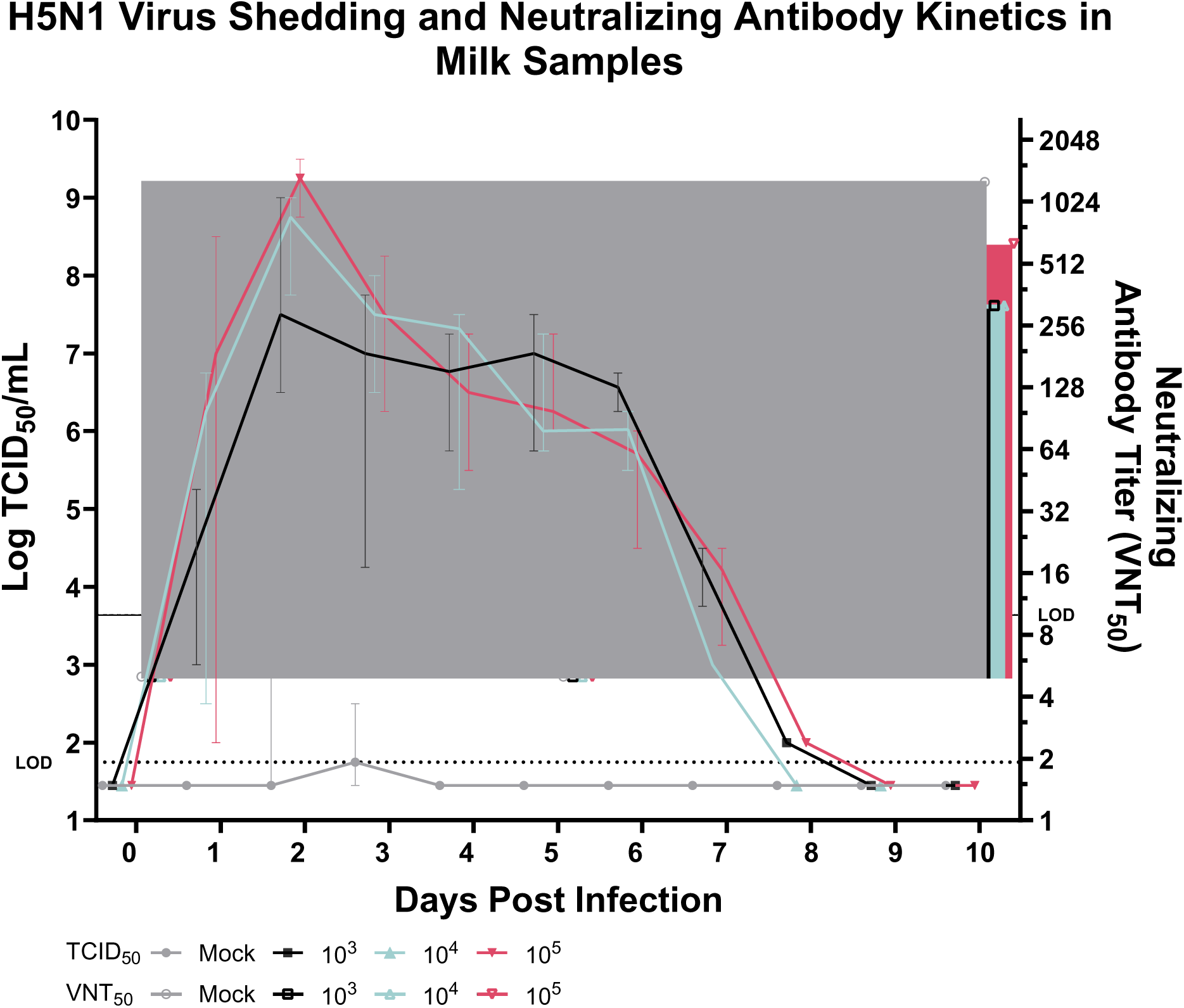
Kinetics of virus shedding and neutralizing antibodies recovered from milk of individual udder quarters. A single plot is used to display the infectious virus titers (left Y-axis) and neutralizing antibody titers (right Y-axis) recovered from milk of individual udder quarters during the study period. Data points represent the median titer of from each udder-quarter; mock (gray circle), low-(black circle), mid-(green triangle), and high-dose (red triangle) inoculated quarters are shown. Error bars represent the range of virus titers recovered from the respective quarters of individual cows at each time point. Neutralizing antibody titers (open symbols) are displayed in the same format. The horizontal dotted-lines represents the limit of detection (LOD) for respective assays, as indicated.

Neutralizing antibodies were detected in both milk and serum of the two cows surviving beyond 5 DPI, beginning at 7 DPI and increasing further by 10 DPI in the last remaining animal (**Supplementary Table 2, 3**). These findings were corroborated by two commercial ELISA kits targeting either the Influenza A nucleoprotein (ID Screen® Influenza A Antibody Competition Multi-Species) or the H5 hemagglutinin (ID Screen® H5 Competition 3.0) (**Supplementary Table 2, 3**). This data confirms seroconversion in both animals surviving beyond 5 DPI.

### Localized Virus Replication in Mammary Tissues

Post-mortem examination and necropsy of individual cows was performed on days 3, 4, 5, 7, and 10 post-infection. An extensive collection of fresh and formalin fixed tissues were sampled across major organ systems (respiratory, gastrointestinal, neurological, and visceral), lymph nodes, individual udder-quarters, and supra mammary lymph nodes. Multiple samples were collected from the udder across distinct anatomical regions: the teat cistern/peri-cistern region and the mammary lobules. Each quarter was processed independently. Tissue homogenates derived from fresh tissue were assayed for IAV-virus RNA and infectious virus (**Supplementary Table 4**). Infectious virus was recovered exclusively from mammary tissues, with only low levels of RNA detected sporadically in other organ systems **(Supplementary Table 4)**.

Cow #89 euthanized on 3 DPI showed high virus titers (10^5^ – 10^7^ TCID_50_/mL) in all H5N1-inoculated udder quarters, with a low but detectable virus titer also present in the mock-inoculated quarter, (**Table 1**). Virus titers in mammary and cisternal tissue were lower (10^3^ – 10^5^ TCID_50_/mL) at 4 DPI (cow #93) and 5 DPI (cow #28), with uninoculated quarters largely negative except for one cisternal sample (cow #28, 5 DPI). In the mammary gland of cow #22 euthanized at 7 DPI, virus was present in the low- and high-dose inoculated quarters, with higher titers in mammary gland sections compared to sections of cisternal tissue. No virus was recovered from mammary gland or cisternal tissues derived from cow #90 euthanized at 10 DPI. In addition, virus was also recovered from supramammary lymph nodes at 3 DPI (10^1^ – 10^3^ TCID_50_/mL), but not at later timepoints (data not shown; RT-qPCR results available in **Supplementary Table 4**).

**Table 1:** Virus titers in mammary gland and cistern.

|  | Udder quarter | 3 DPI | 4 DPI | 5 DPI | 7 DPI | 10 DPI |
| --- | --- | --- | --- | --- | --- | --- |
| **Mammary gland** | Mock | 5.62E+01 | <5.62E+01 | <5.62E+01 | <5.62E+01 | <5.62E+01 |
|  | Low-dose | 3.16E+06 | 1.00E+03 | 1.78E+05 | 1.78E+05 | <5.62E+01 |
|  | Mid-dose | 1.00E+07 | 3.16E+05 | 1.78E+05 | <5.62E+01 | <5.62E+01 |
|  | High-dose | 5.62E+06 | 5.62E+04 | 3.16E+05 | 5.62E+04 | <5.62E+01 |
| **Cistern** | Mock | 5.62E+02 | <5.62E+01 | 5.62E+04 | <5.62E+01 | <5.62E+01 |
|  | Low-dose | 1.00E+05 | 1.00E+04 | 3.16E+04 | 3.16E+01 | <5.62E+01 |
|  | Mid-dose | 3.16E+07 | 1.78E+04 | 1.00E+05 | <5.62E+01 | <5.62E+01 |
|  | High-dose | 5.62E+05 | 3.16E+04 | 5.62E+04 | 1.78E+01 | <5.62E+01 |

*All virus titers are reported as TCID<sub>50</sub>/mL*

### Pathological changes in the mammary gland

#### Gross pathology

At the time of postmortem examination, nearly all cows had visibly evident mastitis predominantly in the right quarters. Externally, affected quarters were moderately enlarged, and warm to the touch. Cutaneous erythema was commonly present in the quarters inoculated with higher doses of virus, and showed moderate to severe edema expanding to the connective tissue fascia in surrounding the udder and in the inguinal and ventral caudal abdominal regions. The swelling and edema extended into the supramammary lymph nodes resulting in moderate to marked enlargement accompanied by medullary congestion and multifocal-petechial hemorrhage of the parenchyma. Each mammary quarter was sectioned from the teat cistern into the mammary lobules after removal of the teat and they were evaluated for enlargement, edema, hyperemia and hemorrhage. The ducts and cisterns were assessed for contents, size and character, color texture, and thickness of the mucosal lining. The gross appearance of the inoculated mammary gland sections was compared to the mammary tissues of the negative control cow (**Figure 3 and Supplemental Figure 4)**. Gross pathological changes became progressively more evident over the duration of the study and are representative of moderate to severe, acute viral mastitis with transition to moderate to severe subacute mastitis by 7-10 DPI. Grossly evident pathological changes were more extensive and severe in the mid- and high-dose inoculated quarters, followed by the low-dose inoculated quarter throughout the entire study; however, by 7 and 10 DPI mild mastitis was also observed in the mock-inoculated quarters. Gross pathological changes observed in the mammary gland at 3 DPI included mild edema and hyperemia of the glandular tissues with slight distention of the teat cistern (**Figure 3A, B**). Similar changes of increased severity, affecting more of the glandular tissue, was seen at 4 DPI with increasing hyperemia of the glandular tissue and the duct mucosa (**Figure 3C**). By 5 DPI, there was moderate to severe ectasia of the mammary cistern and interlobular ducts. The duct mucosa was irregular, nodular and thickened. Lightly adhered clotted milk/fibrin clumps occurred within ducts. Patchy foci of erythema and edema occur throughout the multifocal swollen glandular parenchyma which contained prominent and dilated common lactiferous ducts (**Fig 3D**). Similar and more extensive changes were seen at 7 DPI and were accompanied by enhanced hyperemia and edema of the gland and peripheral connective and adipose tissue. Cistern trabeculae appear turgid, thickened with irregular mottled irregular surface. Common lactiferous ducts are less prominent. Viscous opaque purulent material and clotted milk occlude the duct lumens and lightly adhere to the mucosa of the cistern and ducts. At 10 DPI pathological changes in the udder had progressed. Changes consist of marked ectasia of the teat cistern and lactiferous ducts with partial to complete occlusion of the lumens with tangled mats of thick, firm clotted milk and debris. Mucosal linings were markedly irregular and thickened by numerous multifocal to coalescing nodules while the underlining connective tissue trabecula were expanded and extended into the adjacent parenchyma (edema and fibrosis) (**Figure 3F**). The glandular parenchyma was diffusely edematous and mottled, with pale to light tan regions of erythema. Interlobular ducts were not well visualized. More extensive images of the gross mastitis lesions observed in these cattle are provided in **Supplementary Figure 3** that includes photos of each mammary quarter at the necropsy time points.

**Figure 3:**
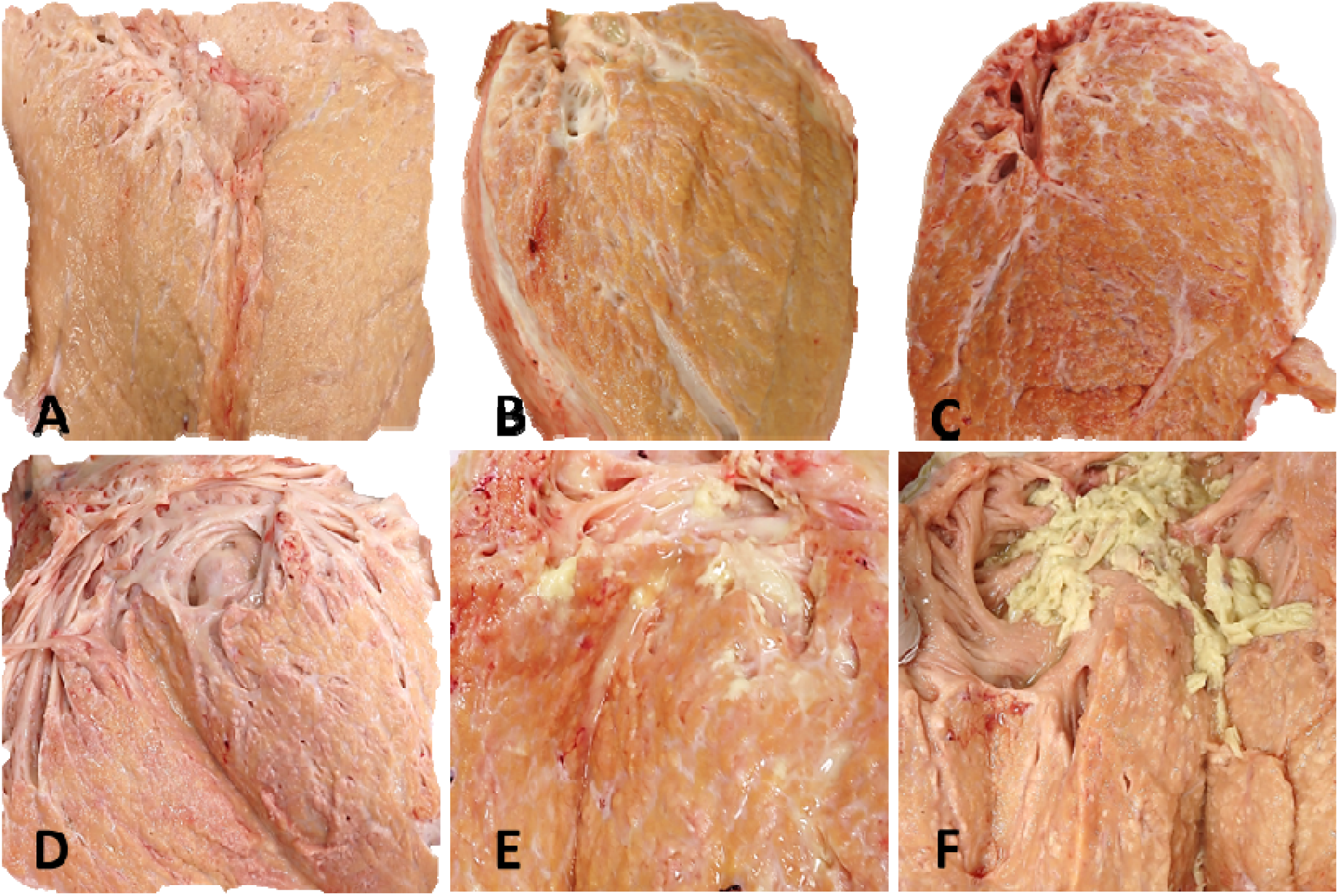
Representative gross pathological changes in the mammary gland following intramammary inoculation. Represented in each panel is **A**) Normal Mammary gland, not inoculated; **B**) 3DPI #89, high-dose (RH). Pathological changes in the mammary glands includes mild edema and hyperemia of the glandular tissue with slight distention of the teat cistern containing water discolored milk. **C**) 4DPI #93, mid-dose (RF). Changes are of increased severity and affect nearly all of the glandular tissue are characterized by increasing hyperemia of the glandular tissues and the duct mucosa. **D**) 5 DPI #28, high-dose (RH). **E**) 7 DPI #22, mid-dose (RF). **F**)10 DPI #90, mid-dose (RF). Acute viral mastitis lesions are mild to moderate at 3-4 DPI (displayed in panel B and C) and become progressively worse over the duration of the study, progressing to moderate to severe subacute mastitis by 7-10 DPI (displayed in panel E and F). On 5 DPI, lesions consist of moderate to severe ectasia of the mammary cistern and interlobular ducts and the duct lining is irregular, nodular and thickened and contains clotted milk. Patchy foci of erythema and edema occur throughout the mammary glandular parenchyma that contains prominent and dilated common lactiferous ducts (D). At 7 DPI, similar although more pronounced changes occur and include glandular hyperemia and marked edema of the gland and peripheral connective tissues. Trabeculae near the cistern are prominent, thickened and have irregular lining. Common lactiferous ducts are less discernable and clotted milk material and fibrin tags occlude lumens (E). At 10 DPI, pathological changes are pronounced and consist of marked ectasia of the cistern and ducts which are partially to completely occluded with thick firm clotted milk and fibrin. Duct linings are markedly irregular and thickened and edema occurs throughout the glandular tissue where interlobular ducts cannot be visualized (E).

#### Histopathology

Extensive histopathology on the mammary tissues of inoculated udders demonstrated acute necrotizing viral mastitis mainly present in the ductular epithelium with variable involvement of the mammary secretory epithelium, dependent of intramammary challenge dose and time of infections. Acute necrotic changes (cattle sampled at 3-5 DPI) gradually changed to more inflammatory, and occasionally suppurative, and reparative responses at later time points (7 and 10 DPI) of the study.

Generally, the mid- and high-dose inoculated quarters exhibited the most prominent and extensive histological changes, relative to normal mammary histology and microanatomy at all collection time points (provided in **Figure 4A**).

**Figure 4:**
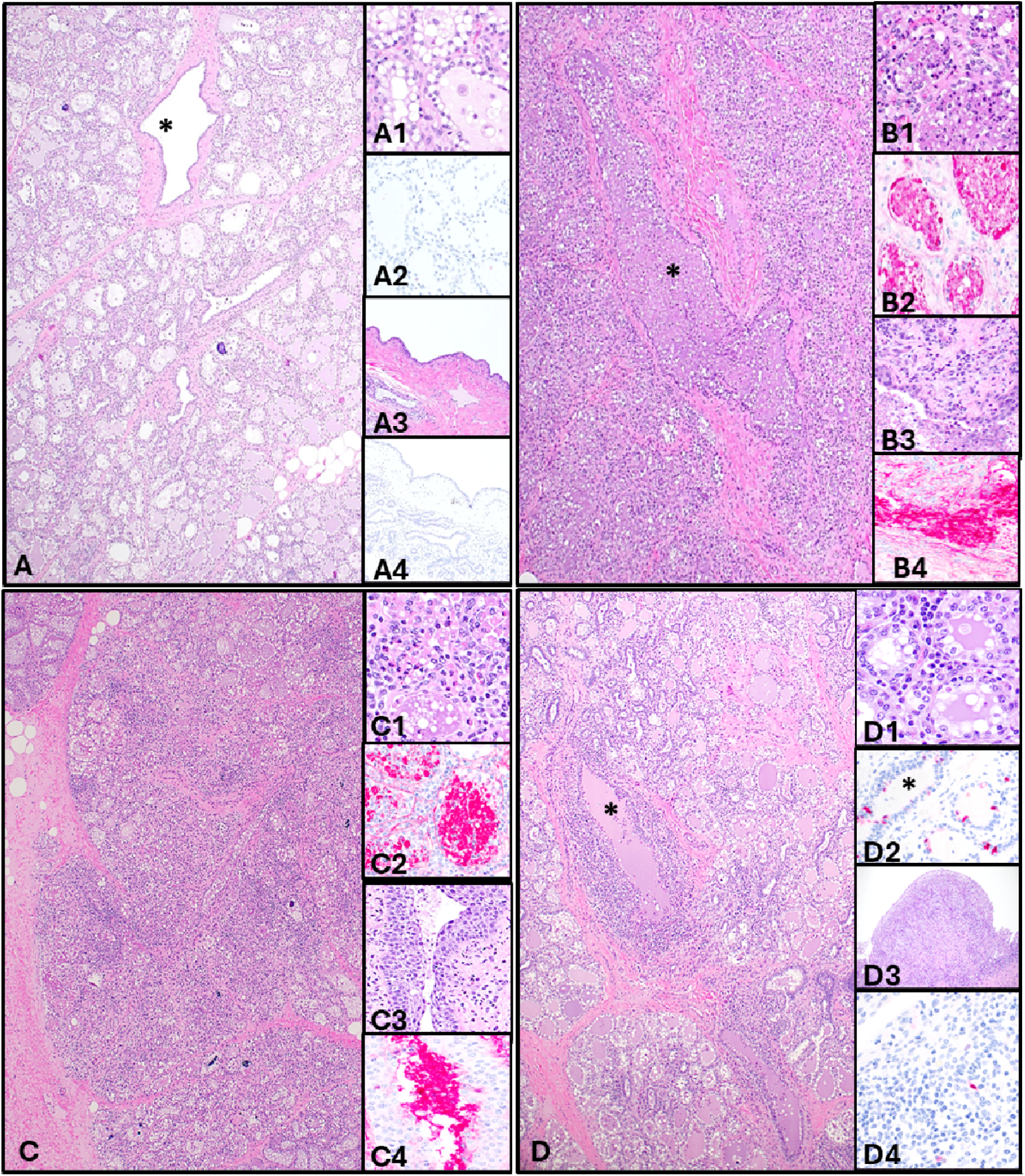
Representative histology and immunohistochemical detection (IHC) of Influenza A antigen in the mammary tissue at 0, 3, 5 and 10 DPI. Intramammary inoculation of individual quarters consisted of either 10^3^ (Left hind, LH), 10^4^ (Right Front; RF), 10^5^ TCID50 (Right hind, RH) of H5N1 clade 2.3.4.4b genotype B1.13, or mock (left front quarter: (LF) uninoculated). (A) Mammary tissue of negative control cow (cow #26, 0 DPI). Normal partially involuted mammary secretory acini are tightly packed and organized in distinct lobules interdigitated with central ducts (*) lined by two layers of low cuboidal epithelium (**A**; 20X: H&E stain). Thick dense connective tissue defines lobules. Secretory acini are suspended in the light vascular interstitium containing loose light infiltrates of lymphocytes and plasma cells. Acini are comprised of variably sized circular to oblong structures surrounded by a single layer of smooth muscle cells. Acini are lined by two layers of epithelium: an inner layer of small flattened basal cells, overlayered with low to high cuboidal secretory epithelium with abundant vacuolated cytoplasm. Acini contain fatty to proteinaceous fluid and infrequently lamellar concretions of mineral or clusters on inflammatory cells (**A1**; 200X: H&E stain). No immunohistochemical detection (IHC) of Influenza A antigen in the non-inoculated mammary tissue (**A2**; 200X: hematoxylin counterstain). Larger common lactiferous ducts and the mammary cistern are line by 2-4 layers of low cuboidal epithelium overlying dense connective tissue, contain small rests of primordial epithelium to variably active acini supported by vascular loose connective tissue (**A3**;100X: H&E stain). No IHC of Influenza A antigen in the non-inoculated mammary tissue (**A4**; 200X: hematoxylin counterstain). (B) Mammary tissue at 3 DPI: High-dose inoculated right hind quarter (cow #89). Glandular tissue architecture is marked disrupted from acinar swelling or collapse and extensive epithelial necrosis of the acinar epithelium and ducts. Edema and mild inflammation widen the supportive stroma and interlobular septa. Ducts are distended by proteinaceous material and necrotic cells (*). The outer layer of ductal epithelium is necrotic or absent; necrosis extends to basement membranes (**B**; 20X: H&E stain). Affected acini are packed with necrotic cellular debris, proteinaceous fluid and fat vacuoles; necrosis can involve the outer secretory layer or both layers leaving exposed basement membrane (**B1**; upper and lower acini; 200X: H&E stain). Intense Congo red positive IHC of Influenza A antigen in necrotic acini contents and epithelia lining when present (**B2**; 200X: hematoxylin counterstain). Larger common lactiferous ducts and the mammary cistern epithelial lining is in various stages of necrosis, hyperplasia, ducts contain large densely cellular aggregates of necrotic cells and large mats of proteinaceous debris and predominately mononuclear inflammatory cells infiltrate the stroma which is edematous (**B3**; 100X: H&E stain). Intense Congo red positive IHC of Influenza A antigen in necrotic epithelia lining a lactiferous duct and its contents (**B4**; 200X: hematoxylin counterstain). (C) Mammary tissue at 5 DPI: Mid-dose inoculated right front quarter (cow #28). Glandular tissue architecture remains moderately disrupted by acinar swelling or collapse and extensive epithelial necrosis of the acini and ducts. Moderate to marked edema and moderate inflammation widen the supportive stroma and interlobular septa. Ducts are distended by proteinaceous material and necrotic cells (*). The outer layer of ductal epithelium is necrotic or absent with basement membranes (**C**; 20X: H&E stain). Shrunken acini contain necrotic cellular debris and increased numbers of mixed inflammatory cells including neutrophils (**C1**; 200X: H&E stain). Intense Congo red positive IHC of Influenza A antigen contents and epithelia lining when present (**C2**; 200X: hematoxylin counterstain). The epithelial lining of larger common lactiferous ducts and the mammary cistern is in various stages of hyperplasia and dysplasia resulting in marked thickening of the epithelial lining; squamous metaplasia and ducts contain aggregates of necrotic cells and proteinaceous debris. Mononuclear inflammatory cells within the edematous stroma (**C3**;100X: H&E stain). Intense Congo red positive IHC of Influenza A antigen contents of the lactiferous duct and only light apical staining of dysplastic epithelium (**C4**; 200X: hematoxylin counterstain). (D) Mammary tissue at 10 DPI: Mid-dose inoculated right front quarter (cow #90). Glandular tissue architecture remains partially disrupted and disorganized. Lobules are variably sized and are separated by increased amounts of edematous and inflamed connective tissue (fibrosis). Acini are irregular sized and lactogenic and while others are shrunken inflamed. Smaller intralobular ducts are hyperplastic, partial ectatic and more numerous, forming into small rest intermixed with inflamed acini or along fibrous planes (*). Larger lactiferous ducts are ectatic, markedly inflamed and distended by proteinaceous material and necrotic cells (*). Moderate to severe inflammation and granulation tissue are thickening the duct wall and extending in to adjacent stroma (**D**; 20X: H&E stain) Shrunken, partially regenerative acini contain proteinaceous material and are lined by large plump partial layers and disorganized hyperplasic low cuboidal epithelium (**D1**; 200X: H&E stain). IHC for Influenza A antigen in acini and hyperplastic glandular ducts demonstrate scattered positive staining in basal epithelial cells (**D2**; 100X: hematoxylin counterstain). The epithelial lining of larger common lactiferous ducts and the mammary cistern is in various stages of hyperplasia and dysplasia, resulting in marked thickening of the epithelial lining, nodule formation and marked squamous metaplasia. Inflammatory cells infiltrate the edematous stroma which is expanded by dense granulation tissue (**D3**; 100X: H&E stain). IHC for Influenza A antigen in these regions demonstrate rare positive staining in basal epithelial cells (**D4**; 200X: hematoxylin counterstain). *\*= Central ducts*

The mid and high -dose inoculated quarter of cow #89 at 3 DPI had extensive disruption of the normal glandular architecture with focally extensive regions of epithelial necrosis of the glandular acini and ductal epithelium (**Figure 4 B, B1, B3**). Nearly all lobules were affected. Glandular acini contained necrotic cellular debris accompanied by mild infiltrates of inflammatory cells within the stroma and acinar lumen. Glandular ducts, lactiferous ducts and ductal lining of the cistern had full to partial thickness, epithelial necrosis and sluffing of necrotic epithelium into the lumen, accompanied by the accumulation of fibrin and pertinacious fluid (**Figure 4, B, B3**). Mild edema expanded interstitial septa and perivascular spaces (**Figure 4, B**). Immunohistochemistry (IHC) for Influenza A viral antigen (nucleoprotein; NP) demonstrated intense antigen accumulation within necrotic acini (**Figure 4,** B2), necrotic ductal epithelium and ductal contents (**Figure 4, B4**). Similar changes as described for 3 DPI were seen in the mid- and high-dose inoculated quarters in cow #93 at 4 DPI.

The acinar and ductular epithelial necrosis remained actively present at 5 DPI in mammary gland samples, however necrotic acini and ducts were commonly lined with a single layer of attenuated basal epithelium (**Figure 4, C, C1**). Affected acini contained necrotic cellular material and/or proteinaceous material accompanied by increased numbers of mixed inflammatory cells including neutrophils. Infrequent mineral deposits (microcalcifications) appeared as concentric layers present in affected and unaffected acini (**Figure 4, C**). The lobular interstitium was irregularly infiltrated by moderate numbers of mononuclear inflammatory cells forming sheets and aggregates often displacing or replacing acini. Moderate to severe edema and early fibrosis thickened the interlobular septa (**Figure 4, C**). Intraglandular ducts in these regions were lined by necrotic or attenuated epithelium, were partially ectatic and contained necrotic cellular debris. In addition, the ductal epithelium in the lactiferous ducts and the cistern was moderately reactive and lined by hyperplastic and dysplastic epithelium of variable thickness forming 10-15 layers of dysplastic partially stratified epithelium (**Figure 4, C3**). Detection for influenza A NP antigen by IHC demonstrated strong positive reaction in necrotic cellular debris in affected secretory acini (**Figure 4, C2**) and positive staining of the lumen contents but not the dysplastic epithelium lining the lactiferous ducts (**Figure 4, C4**). This might indicate continued virus replication primarily in acini. By 7 DPI, mammary samples from cow #22 demonstrated less necrotic changes while hyperplastic and inflammatory changes became more prominent, additionally, there was moderate to severe generalized suppurative mastitis consistent with secondary bacterial infection in the mid-dose inoculated mammary quarter.

At 10 DPI, the final collection day for mammary tissue from cow #90, the glandular changes consisted of mild to moderate interstitial and glandular infiltrates of predominately mononuclear inflammatory cells and fibroblasts as well as supportive collagenous stroma resulting in shrunken, fibrosing, and inflamed lobules (**Figure 4, D**). Affected acini had evidence of regeneration and contained 1-3 layer of pleomorphic epithelium, although some maintained segmental epithelial loss with exposed basement membrane (**Figure 4, D1**). Changes seen in sections at 10 DPI predominately involved the ductular and interstitial compartments of all inoculated quarters. Intralobular ducts moderately distended and were lined by dysplastic stratified squamous epithelium and the duct walls were markedly and variably thickened by inflamed granulation tissue that encroached and partially occluded the duct lumen (**Figure 4, D**) extending out into the duct wall and surround stroma. Changes in the ducts became more pronounced as they increase in size and descend towards the cistern and included more prominent fibrosis and lobule loss. Common lactiferous ducts were circumferentially lined by marked, thickened, and inflamed dysplastic stratified squamous epithelium that commonly formed plaques and nodules, encroaching and partially to completely filling the lumen. The duct lining and nodules were comprised of dysplastic epithelium forming rete pegs that penetrate into the underlying granulation tissue forming the base of the nodule and extending into the markedly thickened wall (**Figure 4, D2**). Numerous lactiferous ducts (LD) were clearly dilated and many segmentally occluded with the inflamed, exuberant granulation tissue covered and infiltrated by dysplastic stratified squamous epithelium. The inflamed granulation tissue partially covered and infiltrated large luminal aggregates of fibrin and degenerate inflammatory cells within the lumen. The wall of affected ducts was thickened by granulation tissue, and early fibrous connective tissue that thickens the surrounding stroma. The mucosal lining of the adjacent cistern was thickened by dysplastic epithelium forming similar nodules of exuberant inflamed granulation tissue that was grossly evident; the reactive tissue extended into the stroma replacing and disrupting fibromuscular components of the udder.

## Discussion

While high-containment work with cattle remains technically demanding, the dry cow model mitigates several of the challenges posed by lactation-associated care and infrastructure requirements. Compared to lactating cows, dry cows require less intensive husbandry, with reduced metabolic requirements (feed intake and diet) and eliminating the need for daily milking, specialized milking systems and continuous disposal of large quantities of infected milk. This reduces facility strain, lowers resource costs, and expands accessibility to institutions with ABSL-3Ag capacity but limited infrastructure for lactation management. Additionally, adult Jersey cows are approximately three-quarters the size of adult Holstein cows, potentially allowing for the inclusion of additional animals in high-containment studies where space constraints and statistical power are limiting factors. Thus, this model offers a practical platform for controlled studies of H5N1 pathogenesis and immune response, particularly in early-stage vaccine and antiviral testing.

This study establishes a novel model using dry dairy (Jersey) cows for investigating highly pathogenic avian influenza virus (HPAIV) H5N1 pathogenesis in cattle. Our findings demonstrate that intramammary infection of dry cows with clade 2.3.4.4b H5N1 (genotype B3.13) results in a clinical disease and pathological course of infection comparable to that observed in lactating cattle, including acute development of severe clinical disease, mastitis, and high-titer virus shedding in milk ^18,20^. As in lactating cows, virus replication (as measured by virus replication) remained primarily localized to the mammary gland, with almost no dissemination to other tissues and organs, and induced a robust neutralizing antibody response detectable in both serum and mammary secretions beyond 7 DPI.

Importantly, this study also provides novel insights into the dose–response relationship for H5N1 intramammary infection and disease progression in infected udder quarters. Quarters inoculated with higher viral doses (mid- and high-dose) exhibited earlier onset and more severe clinical and histological evident mastitis affecting more glandular tissue; this also resulted in a shorter period of time to reach peak viral titers in the milk at 2 DPI as compared to 3-4 DPI in low dose inoculated quarters. By 3 DPI, the high-dose quarters exhibited full thinness necrosis in nearly all ducts of the ductal epithelium with a significant involvement of secretory lobes in the glandular tissue (both in extent of necrosis and numbers of lobules affected); this resulted in large clusters of lobules affected and dispersed through the glandular tissue. In the low-dose inoculated quarters collected at 3 and 4 DPI, there is pronounced virus replication in the ductal epithelium with infrequent clusters of secretory lobes affected which are more generally placed near the cistern or multifocal within the glandular epithelium. Higher virus loads and stronger IHC positivity for the NP antigen was seen at 3 DPI in cisternal and glandular tissues in the mid- and high-dose inoculated quarters when compared to low-dose inoculated quarters; this supports the pathological observations. Low-dose inoculated quarters showed delayed progression to peak virus shedding and generally lower peak virus titers but ultimately reaching similar severity of viral mastitis. Two out of five (40%) low-dose inoculated quarters eventually reached similar peak virus titers as quarters inoculated at higher doses while the remaining 3/5 (60%) peak viral titers remained 2-logs less than observed in higher-dose inoculated quarters.

Mock-inoculated quarters remained largely unaffected, suggesting that the existing anatomical barriers between individual quarters in the bovine udder inhibit viral spread between them. Viral loads in teat swab samples were less consistent, showing high variability between respective quarters across individual study time-points, providing a less accurate representation of virus load in the mammary tissue and in the milk. These findings demonstrate that H5N1 inoculation dose can influence the rate of mastitis progression, its severity, and shedding kinetics, and that the infection remains largely localized to individual udder quarters throughout the early phase of infection. Importantly, these data also confirm the high susceptibility of the bovine udder to H5N1 infection, as inoculation with 10^3^ TCID_50_ was sufficient to establish an active infection, eventually causing severe clinical and pathological outcomes similar to that of quarters inoculated at higher doses.

H5N1-specific antibody responses were detected in all quarters, including those that were mock infected, indicating that systemic antibodies are secreted in all four quarters. This observation supports the idea that a systemic immunization should be able to protect the bovine udder, provided the correct type and magnitude of vaccine response is sufficient for protection.

Despite its strengths, the dry cow model has inherent limitations. Milk production during dry-off is minimal, or possibly even absent, making this model less suited for transmission studies, particularly where exposure via milk handling or milking equipment is believed to play a central role ^8,11,18^. Accordingly, this platform is best suited for studying within-host dynamics, mammary pathogenesis, and immune responses, rather than evaluating horizontal transmission pathways.

In summary, the dry cow model described here recapitulates key features of HPAIV H5N1 clade 2.3.4.4b genotype B3.13 infection observed in lactating cattle and offers a more scalable, cost-effective alternative for studying bovine H5N1 pathogenesis and immunity in high-containment livestock research settings. These results also support a current hypothesis suggesting that milk and milk-associated activities remain central to H5N1 transmission dynamics, as detections of HPAIV H5N1 and severe clinical disease in cows during the dry phase are reported less often in the field. Our observations also suggest that H5N1 influenza virus replication in the bovine udder is compartmentalized to a degree not previously appreciated, with important implications for pathogenesis and within-host viral evolution.

## Materials & Methods

### Ethics Statement / Biosafety

All experiments were conducted at Kansas State University and approved by the Kansas State University (KSU) Institutional Biosafety Committee (IBC, Protocol # 1758) and the Institutional Animal Care and Use Committee (IACUC, Protocol # 4992) in compliance with the Animal Welfare Act. All experiments involving infectious highly pathogenic avian influenza virus were conducted in biosafety level-3+ (BSL-3+) and/or animal biosafety level 3-agriculture (ABSL-3Ag) at the Biosecurity Research Institute (BRI) at KSU in Manhattan, KS, USA. Select samples were subjected to validated inactivation protocols/methods prior to transfer to lower containment levels for further evaluation, where enhanced biosafety level practices were implemented.

### Virus and Cells

The North American HPAIV H5N1 isolate A/Cattle/Texas/063224-24-1/2024, genotype B3.13 (GIS AID accession number: EPI_ISL_19155861) was used for inoculation of cattle and virus neutralizing antibody assays titers in serum and milk. Bovine uterine epithelial cells (CAL-1; In-house) were used for initial isolation and propagation of virus stocks. One additional passage on Madin-Darby canine kidney (MDCK) cells was used to propagate the virus stock for applications in the present experiment. The virus stock used for inoculation was subjected to next generation sequencing workflows and displayed high sequence identity with the original virus isolate. Virus titrations and neutralization assays were also evaluated on MDCK cells. Cultures were maintained in Dulbecco’s Modified Eagle Medium (DMEM; Corning, Manassas, VA, USA), supplemented with 5% fetal bovine serum (FBS; R&D systems, Flower Branch, GA, USA) and 1% antibiotic-antimycotic solution (Gibco, Grand Island, NY, USA). For virus cultivation and assays, media was deprived of FBS and supplemented with 0.3% bovine serum albumin (BSA; Sigma-Aldrich, Darmstadt, Germany) and 1% minimum essential medium vitamin solution (Gibco, Grand Island, NY, USA), in addition to 1 µg/mL of TPCK-treated trypsin.

### Experimental Design

#### Pre-inoculation: Screening and baseline clinical evaluation

Six female Jersey (*Bos taurus taurus*) cows three days into the dry-off period following first lactation (mono-parous) were transported from a Nebraska livestock operation to Kansas State University College of Veterinary Medicine (KSU-CVM) in Manhattan, KS. Cattle were housed in open outdoor penning for 14-days post-arrival for acclimation and transition to new dietary rations. Cattle were relocated to a secure ABSL-3Ag facility nearby for the remainder of the study. Prior to inoculation, individual baseline parameters were determined for feed intake (qualitative measure), rectal temperature, behavior, and other criteria. Clinical samples (swabs and blood) were also collected and screened for IAV-specific RNA and antibodies as well as ten common bovine respiratory disease pathogens (**Supplementary Table 1**). Milk samples were collected from individual quarters and evaluated (semi-quantitively) to determine baseline somatic cell counts using the California Mastitis Test (CMT) and retained for virus titration and immunoassay.

#### Intramammary inoculation of udder quarters with escalating titers of HPAIV H5N1

Jersey cows were inoculated intramammary with clade 2.3.4.4b HPAIV H5N1 (genotype B3.13) at day 21 of the dry-off period. In preparation for intramammary inoculation, teats were thoroughly cleansed with water to remove debris and disinfected with alcohol. Keratin plugs, if present, were dislodged as residual milk was stripped from each quarter. Syringes, affixed with plastic 3.125” udder infusion cannulas (Jorgensen Laboratories, SKU J0277CP; Colorado, USA) were used to administer 0.5mL of inoculum intramammary. Udder quarters were designated as mock-inoculated (left front quarter), low-dose inoculated (left hind quarter), mid-dose inoculated (right front quarter), and high-dose inoculated (right hind quarter); administered 0.5mL of DMEM only or HPAIV H5N1 inoculum at 10^3^ TCID_50_, 10^4^ TCID_50_, or 10^5^ TCID_50_, respectively (**Supplementary Figure 1**). As cannulas were removed, gentle pressure was applied by hand to clinch the teat cistern for 30-60seconds and occlude immediate evacuation of inoculum. Iodine was applied to each teat after inoculation.

#### Post-inoculation: Clinical evaluations, sampling schedule, sequential necropsy

Thorough examinations were conducted daily by licensed veterinarians to document overt clinical symptoms, evaluate udder quarters for mastitis, and monitor changes in rectal temperature and feed intake **(Supplementary Table 6).** Although cows were in the dry-off period and did not require daily milking, small volumes of milk were still produced and collected throughout the study period.

Regular time-point collections of clinical samples and milk were used to assess the route and kinetics of virus shedding (**Supplementary Figure 1**). Swabs were collected in 2 mL of DMEM containing 1% antibiotic/antimycotic solution. Collection time-points for swabs were as follows: nasal-, oral-, rectal-, and teat swabs (-1, 1-10 DPI), vaginal swabs (4-10 DPI), whole blood (-1, 1-7, 10 DPI), serum (-1, 5, 7, 10 DPI). Milk from individual udder quarters was also collected from 0-10 DPI. Working sequentially from mock-inoculated to high-dose inoculated quarters, each teat was cleaned of any gross debris with water and disinfected with alcohol wipes before gently stripping milk by hand into individual collection vessels. A portion of milk was immediately used for semi-quantitative determination of somatic cell counts using the California Mastitis Test (CMT), per manufacturer instructions (Jorgensen Laboratories, SKU J037OK; Colorado USA). Following collection, teats were cleaned with iodine-soaked gauze pads and personnel changed gloves before proceeding with sample collection.

To determine the scope and extent of IAV infection and virus dissemination, post-mortem examinations were conducted at 3, 4, 5, 7, and 10 DPI (*n=1/day)*. The udder was removed in-toto and set apart from the carcass as sampling collection and processing was conducted. All remaining tissues were observed in-situ and apparent gross lesions were documented prior to extensive paired sample collection of fresh tissues and for formalin fixation. Extensive postmortem evaluation of the mammary gland and reproductive system were conducted along with tissues from all multiple organ systems including the upper and lower respiratory tract visceral organs, lymphoid, gastrointestinal, ocular and central nervous and urogenital system. Fresh tissues were later processed into homogenates for virus isolation and molecular testing. Tissues for histology were processed into 10% neutral buffered formalin for 5 days then transferred to 70% ethanol then trimmed and embedded in paraffin following standard tissue processing in the histology laboratory for the Kansas State Veterinary Diagnostic Laboratory (KSVDL) for histological evaluation of hematoxylin and eosin-stained slides and IAV-specific IHC for nucleoprotein antigen following standard protocols at the KSVDL. Briefly Influenza A IHC uses a 1:4000 dilution of the H1N1 NP rabbit antibody (GenScript) in a 15-minute incubation following a 10-minute,100°C, high PH EDTA antigen recovery protocol and the Bond Polymer Refine Red Detection kit using a 10 minute fast red chromogen, and counterstain with hematoxylin.

### Nucleic acid extraction and (IAV-specific/targeted) RT-qPCR

Nucleic acid was extracted from clinical samples, tissue homogenates (10% weight:volume), and whole milk samples according to previously established protocols ^18^. Briefly, 200µL of sample lysate, containing an equal volume of liquid sample and RLT lysis buffer (Qiagen), was used for extraction with an automated magnetic bead-based extraction system (Taco Mini, GeneReach; BioSprint96, Qiagen) using commercial reagents (GeneReach).

Amplification and relative quantification of IAV RNA was determined using a one-step RT– qPCR assay targeting the matrix gene segment of IAV employing a modified M + 64 probe. Samples were tested in duplicate wells and cycle thresholds (Ct) were determined using the Bio-Rad CFX96 Optics Module C1000 Touch (Bio-Rad) with thermocycling conditions as described previously ^7^. The limit of detection for this RT-qPCR assay was determined from a standard curve generated from 10-fold serial dilutions of viral RNA of known concentration. Samples were considered *i)* positive when both reactions resulted in Ct-values < 38, *ii)* suspect-positive if one of two reactions were positive, or *iii)* negative if one or both reactions were above 38 Ct.

### Virus titration and IAV-neutralizing antibody assays

Infectious virus titers were determined using previously established methods for virus titration and immunofluorescence assay (IFA) on Marian-derby canine kidney cells^33^. Virus titers were determined for all milk samples, mammary-associated tissues (cistern, mammary gland, super-mammary lymph nodes), and any clinical samples or tissues with Ct<28. Briefly, whole milk and tissue homogenates were clarified by centrifugation and filtered through a 0.45µm syringe filter prior to titration. Samples were titrated in four replicates on 96-well plates using 10-fold serial dilutions and transferred onto 96-well plates containing confluent monolayers of MDCK cells. Following 48 hours of incubation at 37°C, cells were fixed with ice-cold methanol and incubated with the HB65 monoclonal antibody targeting the IAV nucleoprotein ^34^. After washing, cells were incubated with anti-mouse antibodies conjugated with Alexa Fluor 488 (Invitrogen, Carlsbad, CA, USA). Virus titers were calculated using the Reed-Muench method ^35^.

### IAV-specific antibody detection

Serum and whole milk samples were evaluated for the presence of IAV nucleoprotein and H5-subtype specific antibodies using commercial competitive enzyme-linked immunoassays (ID Screen® Influenza H5 Antibody Competition 3.0 Multi-species Version 3; ID Screen® Influenza A Antibody Competition Multi-Species; Innovative Diagnostics) according to manufacturer’s instructions for respective bovine biologicals.

IAV-specific neutralizing antibody titters were also determined in serum and milk, using previously established protocols with slight modifications ^18,36^. In brief, serum and whey fractions of whole milk, clarified by centrifugation, were diluted (1:10) and heat inactivated at 56°C for 30 minutes, then combined with an equal volume of H5N1 B3.13 stock virus, diluted to 100 TCID_50_ per 50 µl, in duplicate wells on 96-well plates and incubated at 37 °C for 1 h. Subsequently, the serum/virus mixture was transferred to 96-well plates containing confluent monolayers of MDCK cells. After 48 hours of incubation, cells were fixed with methanol, and IFA was performed as described above for virus titration. Neutralizing antibody titters were recorded as the final dilution at which 50% inhibition of virus growth was observed per well.

## Acknowledgments

Invaluable contributions that supported the success of this work were made by the professional and technical associates and staff of KSU-CEEZAD/CEZID personnel, including M. Ardalan, S. Ghimire, P. Assato, D. Madden, Y. Li, and Z. Kohl. Additional support was provided by KSU-VDL molecular diagnostic and histopathology laboratory personnel and leadership (J. Retallick, G. Hazlicek and L. Knoll). We also thank the KSU Comparative Animal Group and support staff of the Biosecurity Research Institute (BRI) at Kansas State University for their coordination and oversight. We thank David O’Connor and Nicholas Minor of the University of Wisconsin School of Medicine and Public Health for helpful discussions.

## Author Contributions

Conceptualization: J.A.R., D.G.D., I.M., N.N.G., K.C., J.D.T, T.C.F.; Data curation: K.C., J.D.T., G.S., S.E., E.L., F.M.F., N.N.G., M.C., W.W., H.M.M., T.C.F., B.C.; Methodology: K.C., J.D.T, T.K., E.L., S.K., C.D.M., W.C.W., S.E., M.C., D.G.D., F.M.F., W.W., H.M.M., T.C.F., B.C.; Formal analysis: J.A.R., K.C., J.D.T., W.W., H.M.M., T.C.F., B.C.; Investigation: J.A.R., I.M., K.C., J.D.T., T.K., G.S., S.K., C.D.M., S.E., E.L., G.V., F.M.F., W.C.W., N.N.G., W.W., H.M.M., T.C.F., M.C., D.G.D.; Visualization: K.C., J.D.T., N.N.G., W.W., H.M.M.; Writing, original draft: K.C., J.D.T.; Writing, review and editing: J.A.R., J.D.T., K.C., T.K., G.S., G.V., F.M.F., W.C.W., N.N.G., D.G.D., W.W., H.M.M., T.C.F., B.C.; Supervision: J.A.R., I.M., K.C., J.D.T., T.C.F.; Funding acquisition: J.A.R., D.G.D., N.N.G., I.M., T.C.F., H.M.M.

