## Supplemental Material for "Pathogenesis of H5N1 Clade 2.3.4.4b in dry Jersey cows following intramammary inoculation shows within-host compartmentalization"

### Supplementary Figures and Tables

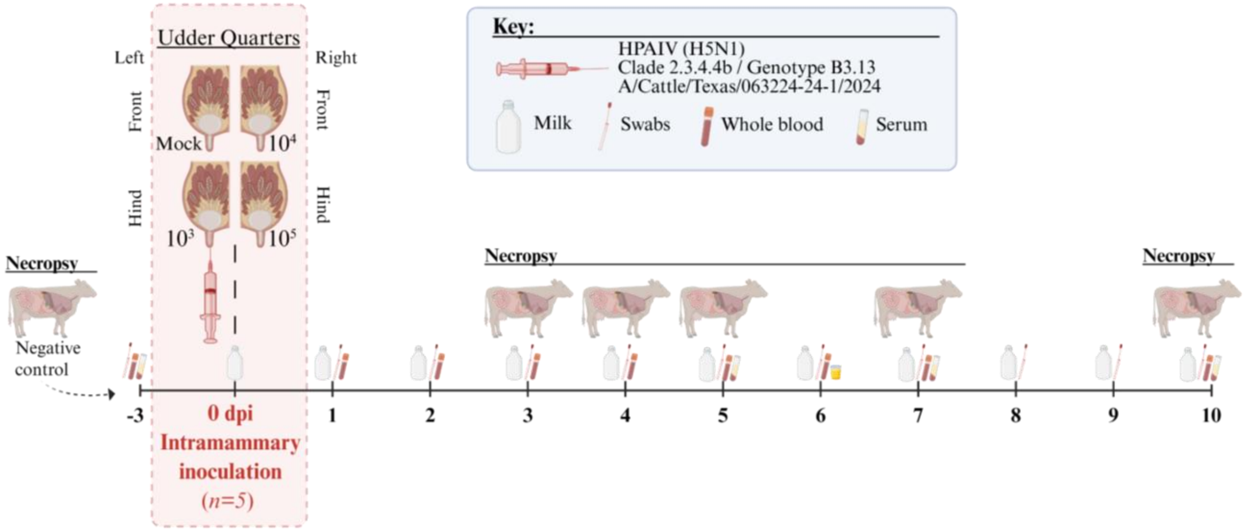


**Supplementary Figure 1: Experimental Design.** A schematic representation of the intramammary HPAIV challenge, sample collection, and sequential necropsy schedule is shown. Five monoparous Jersey cows, 21-days into the dry period, were inoculated intramammary with HPAIV H5N1 clade 2.3.4.4b (genotype B3.13). To determine the impact of viral dose on clinical and pathological outcomes, individual quarters for each cow were inoculated with $10^3$, $10^4$ or $10^5$ $TCID_{50}$ per quarter with one left as a mock-inoculated control. Clinical observations and sampling were conducted on a daily basis to monitor overall health conditions and viral shedding. Post-mortem examinations were conducted on days 3, 4, 5, 7, and 10 post-infection (n=1/day). Created in Biorender.(https://BioRender.com).

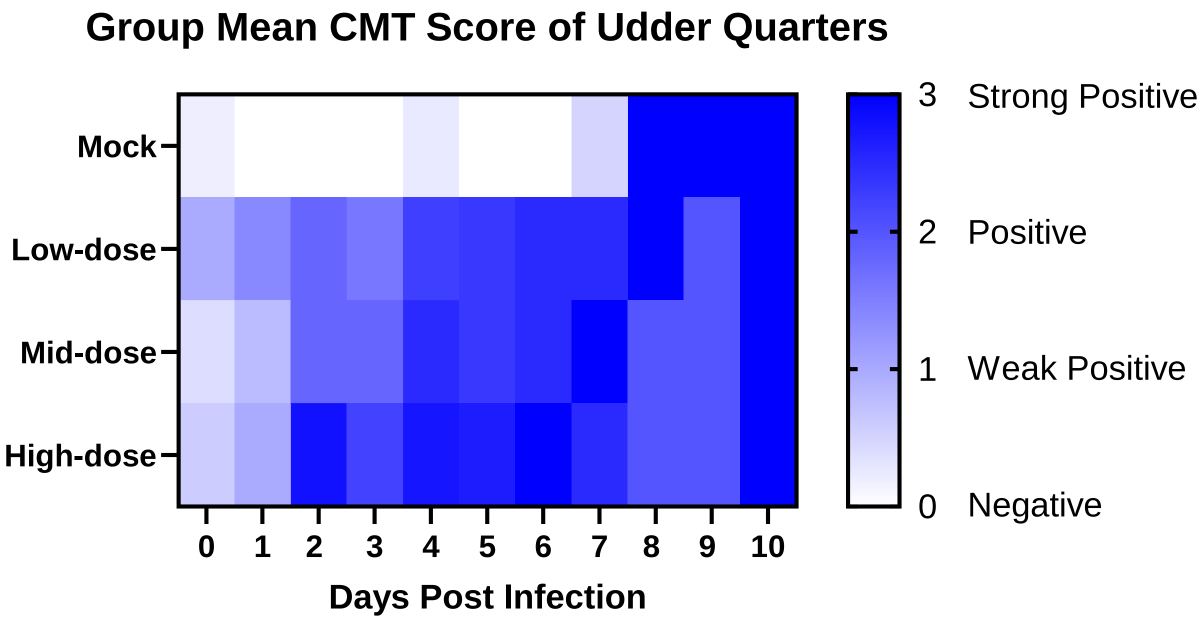


**Supplementary Figure 2: Results of California Mastitis Test (CMT) presented as a heat-map.** CMT scores (0-3; Negative-Strong positive) from udder quarters were grouped according to the dose of inoculum administered at 0 DPI (mock-inoculated; low-dose, $10^3$; mid-dose, $10^4$; or high-dose, $10^5$ $TCID_{50}$ in 0.5mL per quarter) and the grouped-mean CMT score from respective quarters on each day post-infection is displayed. Dark-blue hues represent high mean CMT scores from udder quarters inoculated at respective doses on each day, light-blue hues indicate lower CMT scores.

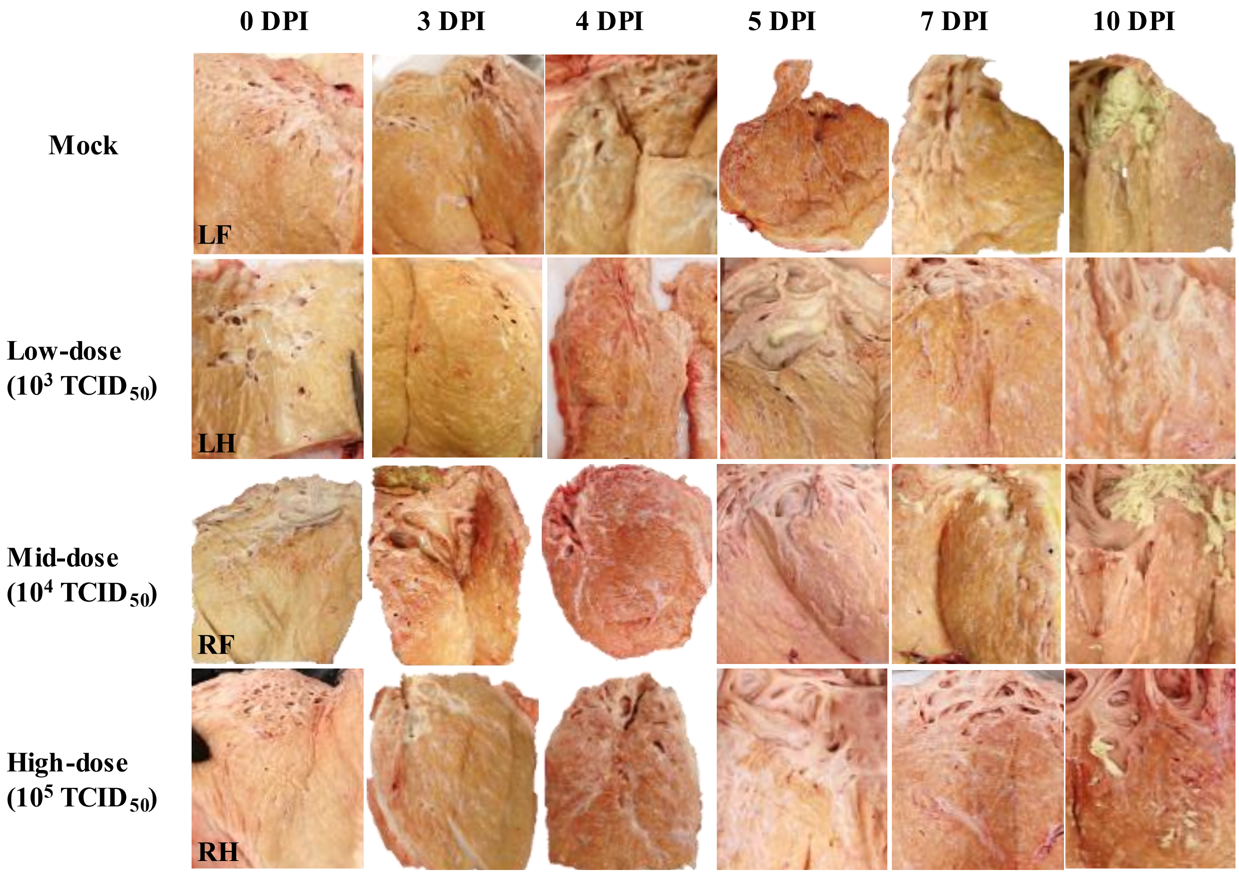


**Supplemental Figure 3: Gross mastitis lesions in mammary quarters at necropsy time-points.** Gross mastitis lesions from individual udder quarters are displayed for each cow at respective necropsy time-points. Apparent changes in the severity of mastitis can be appreciated between quarters that were mock inoculated (top), and quarters that were inoculated with low-, mid-, or high-dose (bottom) of H5N1 inoculum. The progression of mastitis and tissue damage can also be observed from representative sections collected from normal tissue, collected from the negative control cow at 0 DPI (left), and those collected at 3-4 DPI, 5 DPI, and 7-10 DPI (right). Clotted and discolored milk can also be observed in all sections collected at 10 DPI.

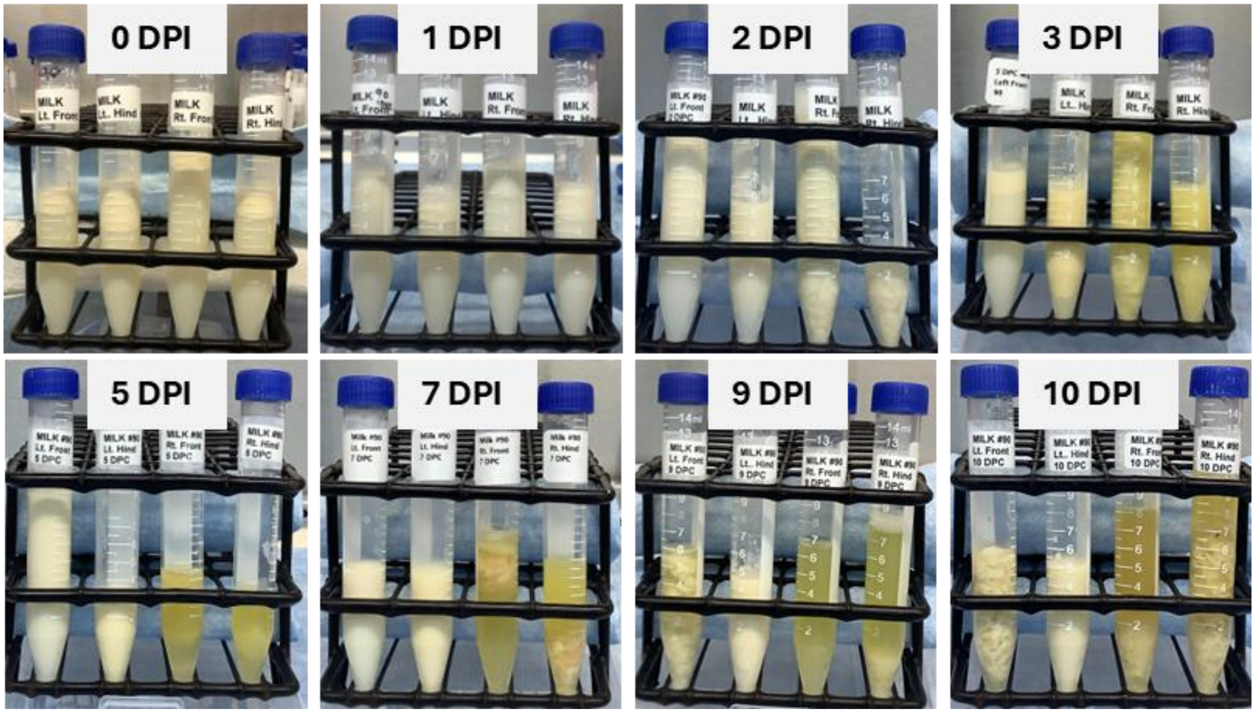


**Supplemental Figure 4: Reduction in milk quality over time and between udder quarters inoculated with different virus titers.** Milk samples collected from individual quarters that were inoculated with increasing titers of HPAIV H5N1 are depicted. Samples in each image represent a single time-point collection and are arranged from mock inoculated (left), to low-, mid-, and high-dose inoculated.

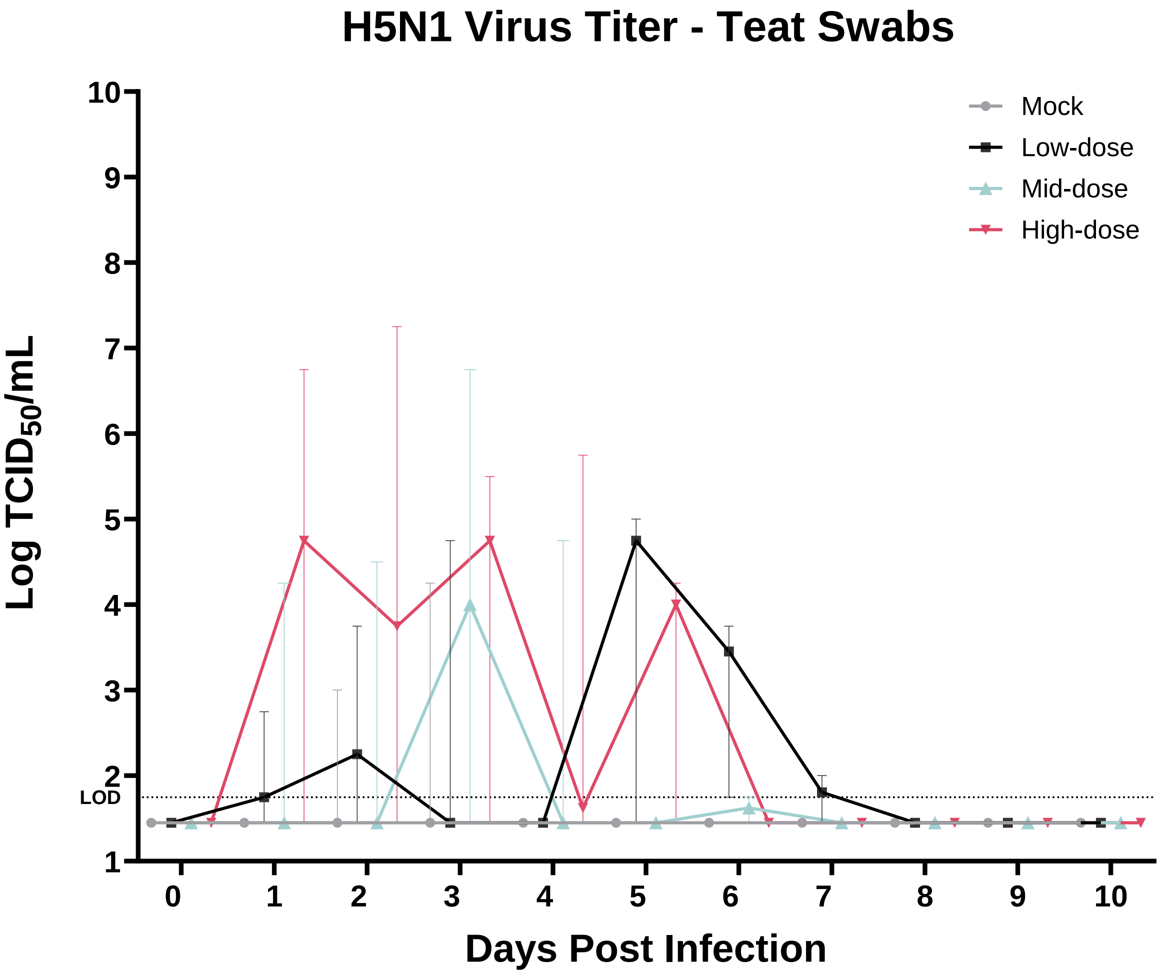


**Supplementary Figure 5: IAV-Matrix gene detection in teat swabs.** External teat swabs were collected daily from each udder quarter to evaluate H5N1 virus shedding. H5N1 virus titers are shown. Colors correspond to udder quarters that were either mock inoculated or inoculated with low-, mid-, or high-dose of H5N1. The limit of detection for infectious virus is shown as a dotted line. Individual data points represent the mean titer of virus detected in respective swabs from all udder quarters of all cows at each timepoint. Error bars show the standard deviation.

**Supplementary Table 1: Bovine Respiratory Disease PCR Panel.**

| Cow ID | Bovine Viral Diarrhea Virus | Bovine Herpesvirus 1 | Bovine Respiratory Syncytial Virus | Mycobacterium bovis | Bovine Coronavirus |
|---|---|---|---|---|---|
| 93 | Negative | Negative | Negative | Negative | Negative |
| 22 | Negative | Negative | Negative | Negative | Negative |
| 90 | Negative | Negative | Negative | Negative | Negative |
| 28 | Negative | Negative | Negative | Negative | Negative |
| 89 | Negative | Negative | Negative | Negative | Negative |
| 26 | Negative | Negative | Negative | Negative | Negative |
| 26* | Negative | Negative | Negative | 22.76 | Negative |

| Cow ID | Influenza D Virus | Mannheimia haemolytica | Pasteurella multocida | Histophilus somni | Bibersteinia trehalosi |
|---|---|---|---|---|---|
| 93 | Negative | Negative | Negative | Negative | Negative |
| 22 | Negative | Negative | Negative | Negative | Negative |
| 90 | Negative | Negative | Negative | Negative | Negative |
| 28 | Negative | Negative | Negative | Negative | Negative |
| 89 | Negative | 37.08 | 33.26 | Negative | Negative |
| 26 | Negative | Negative | Negative | Negative | Negative |
| 26* | Negative | Negative | 25.17 | Negative | Negative |

Interpretation of results: Positive = Ct values <36; Suspect/Inconclusive = Ct values between 36 and 39; Negative = Ct values > 39 or 0. * Lung Abscess

### Supplementary Table 2: IAV-specific and neutralizing antibodies in milk.

| | | | Day Post-Infection | | | | | | |
|---|---|---|---|---|---|---|---|---|---|
| Cow ID | Udder quarter | ASSAY | -3 | 0 | 5 | 7 | 8 | 9 | 10 |
| 26 | Pooled | $VNT_{50}$ | <1:10 | | | | | | |
| | | H5 | - | | | | | | |
| | | NP | - | | | | | | |
| 89 | L/F | $VNT_{50}$ | | <1:10 | | | | | |
| | | H5 | | - | | | | | |
| | | NP | | - | | | | | |
| | L/H | $VNT_{50}$ | | <1:10 | | | | | |
| | | H5 | | - | | | | | |
| | | NP | | - | | | | | |
| | R/F | $VNT_{50}$ | | <1:10 | | | | | |
| | | H5 | | - | | | | | |
| | | NP | | - | | | | | |
| | R/H | $VNT_{50}$ | | <1:10 | | | | | |
| | | H5 | | - | | | | | |
| | | NP | | - | | | | | |
| 28 | L/F | $VNT_{50}$ | | <1:10 | <1:10 | | | | |
| | | H5 | | - | - | | | | |
| | | NP | | - | - | | | | |
| | L/H | $VNT_{50}$ | | <1:10 | <1:10 | | | | |
| | | H5 | | - | - | | | | |
| | | NP | | - | - | | | | |
| | R/F | $VNT_{50}$ | | <1:10 | <1:10 | | | | |
| | | H5 | | - | - | | | | |
| | | NP | | - | - | | | | |
| | R/H | $VNT_{50}$ | | <1:10 | <1:10 | | | | |
| | | H5 | | - | - | | | | |
| | | NP | | - | - | | | | |
| 90 | L/F | $VNT_{50}$ | | <1:10 | <1:10 | <1:10 | 1:40 | 1:160 | 1:1280 |
| | | H5 | | - | - | + | + | + | + |
| | | NP | | - | - | + | + | + | + |
| | L/H | $VNT_{50}$ | | <1:10 | <1:10 | 1:10 | 1:20 | 1:160 | 1:320 |
| | | H5 | | - | - | - | + | + | + |
| | | NP | | - | - | - | - | + | + |
| | R/F | $VNT_{50}$ | | <1:10 | <1:10 | 1:40 | 1:40 | 1:160 | 1:320 |
| | | H5 | | - | - | + | + | + | + |
| | | NP | | - | - | - | + | + | + |
| | R/H | $VNT_{50}$ | | <1:10 | <1:10 | 1:20 | 1:160 | 1:320 | 1:640 |
| | | H5 | | - | - | -/+ | + | + | + |
| | | NP | | - | - | - | + | + | + |
| 22 | L/F | $VNT_{50}$ | | <1:10 | <1:10 | 1:20 | | | |
| | | H5 | | - | - | - | | | |
| | | NP | | - | - | + | | | |
| | L/H | $VNT_{50}$ | | <1:10 | <1:10 | 1:40 | | | |
| | | H5 | | - | - | - | | | |
| | | NP | | - | - | - | | | |
| | R/F | $VNT_{50}$ | | <1:10 | <1:10 | 1:20 | | | |
| | | H5 | | - | - | -/+ | | | |
| | | NP | | - | - | + | | | |
| | R/H | $VNT_{50}$ | | <1:10 | <1:10 | 1:40 | | | |
| | | H5 | | - | - | - | | | |
| | | NP | | - | - | - | | | |
| 93 | L/F | $VNT_{50}$ | | <1:10 | | | | | |
| | | H5 | | - | | | | | |
| | | NP | | - | | | | | |
| | L/H | $VNT_{50}$ | | <1:10 | | | | | |
| | | H5 | | - | | | | | |
| | | NP | | - | | | | | |
| | R/F | $VNT_{50}$ | | <1:10 | | | | | |
| | | H5 | | - | | | | | |
| | | NP | | - | | | | | |
| | R/H | $VNT_{50}$ | | <1:10 | | | | | |
| | | H5 | | - | | | | | |
| | | NP | | - | | | | | |

*Interpretation: (-) = negative; (-/+) = doubtful; (+) = positive*
*Shaded cells indicate that samples were not available for testing*

**Supplementary Table 3: IAV-specific and neutralizing antibodies in serum.**

| | | Day Post-Infection | | | | | | |
|---|---|---|---|---|---|---|---|---|
| Cow ID | ASSAY | -3 | -1 | 3 | 4 | 5 | 7 | 10 |
| 26 | $VNT_{50}$ | <1:10 | | | | | | |
| | H5 | - | | | | | | |
| | NP | - | | | | | | |
| 89 | $VNT_{50}$ | | <1:10 | <1:10 | | | | |
| | H5 | | - | - | | | | |
| | NP | | - | - | | | | |
| 28 | $VNT_{50}$ | | <1:10 | | | <1:10 | | |
| | H5 | | - | | | - | | |
| | NP | | - | | | - | | |
| 90 | $VNT_{50}$ | | <1:10 | | | <1:10 | 1:40 | 1:640 |
| | H5 | | - | | | - | + | + |
| | NP | | - | | | - | + | + |
| 22 | $VNT_{50}$ | | <1:10 | | | <1:10 | 1:40 | |
| | H5 | | - | | | - | - | |
| | NP | | - | | | - | + | |
| 93 | $VNT_{50}$ | | <1:10 | | <1:10 | | | |
| | H5 | | - | | - | | | |
| | NP | | - | | - | | | |

*Interpretation: (-) = negative; (-/+) = doubtful; (+) = positive*
*Shaded cells indicate that samples were not available for testing*

**Supplementary Table 4: H5-subtype specific RNA in tissues.**

| Tissue: Post-mortem: / Cow ID: | 3 DPI<br>89 | 4 DPI<br>93 | 5 DPI<br>28 | 7 DPI<br>22 | 10 DPI<br>90 |
|---|---|---|---|---|---|
| Eye lid | - | - | - | - | - |
| Third eye lid | / | - | - | - | - |
| Nasal concha | / | - | - | - | - |
| Ethmoturbinates | - | - | - | - | - |
| Brain | - | - | - | - | - |
| Olfactory Bulb | - | - | - | - | - |
| Treachea (Pool) | - | - | - | - | - |
| Bronchi | / | - | - | - | - |
| Lung | - | - | - | - | - |
| Lung lesion | - | - | - | - | - |
| Heart | - | - | - | - | - |
| Liver | - | - | - | - | - |
| Kidney | - | - | - | - | - |
| Spleen | - | - | - | - | - |
| Pancreas | / | - | - | - | - |
| Gastrointestinal (Pool) | - | - | - | - | - |
| Feces | - | - | - | - | - |
| Uterus | - | - | - | - | - |
| Vagina | + | - | - | - | - |
| Placenta | x | x | - | x | x |
| Thymus | - | - | - | - | - |
| Tonsil | - | - | - | - | - |
| Pharyngeal tonsil | + | - | - | - | - |
| Retropharyngeal LN | - | - | - | - | - |
| Mandibular LN | - | - | - | - | - |
| Cranial mediastinal LN | - | - | - | - | - |
| Tracheobronchial LN | - | - | - | - | - |
| Ileo-cecal LN | - | - | - | - | x |
| Superficial cervical LN | - | - | - | - | - |
| Mesentaric LN | - | - | - | - | - |
| Gastrohepatic LN | - | - | - | - | - |
| Right Prefemoral LN | - | - | - | - | - |
| Left Prefemoral LN | - | - | - | - | - |
| Retroperitoneal LN | - | - | - | - | - |
| Super Mammary LN 1 | + | x | x | + | x |
| Super Mammary LN 2 | + | - | - | - | x |
| Super Mammary LN 3 | + | - | ++ | - | - |
| Super Mammary LN 4 | ++ | + | x | x | x |
| Super Mammary LN 5 | ++ | + | ++ | - | x |
| Super Mammary LN 6 | + | + | x | + | - |
| Left Axillary LN | - | - | - | - | - |
| Right Axillary LN | - | - | - | - | - |

Abbreviations*: LN = Lymph node*

Symbols*: (x) = Not collected; (-) = Not detected; (/) = Suspect-positive; (+) = Positive, 38>Ct>31; (++) = Strong positive, Ct<31*

**Supplementary Table 5: H5-subtype specific RNA detection in clinical samples.**

| | Cow ID | Day-post infection -1 | 1 | 2 | 3 | 4 | 5 | 6 | 7 | 8 | 9 | 10 |
|---|---|---|---|---|---|---|---|---|---|---|---|---|
| Nasal | 89 | - | - | + | + | X | X | X | X | X | X | X |
| | 28 | - | - | + | + | - | - | X | X | X | X | X |
| | 90 | - | - | + | / | + | - | / | - | + | - | - |
| | 22 | - | - | ++ | + | + | + | + | - | X | X | X |
| | 93 | - | - | + | - | - | X | X | X | X | X | X |
| | 26 | - | X | X | X | X | X | X | X | X | X | X |
| Oral | 89 | - | - | + | - | X | X | X | X | X | X | X |
| | 28 | - | - | + | + | - | - | X | X | X | X | X |
| | 90 | - | - | - | - | + | + | + | / | - | + | - |
| | 22 | - | - | ++ | + | - | + | / | - | X | X | X |
| | 93 | - | - | + | / | - | X | X | X | X | X | X |
| | 26 | - | X | X | X | X | X | X | X | X | X | X |
| Rectal | 89 | - | - | / | + | X | X | X | X | X | X | X |
| | 28 | - | - | + | + | - | - | X | X | X | X | X |
| | 90 | - | - | - | - | + | + | + | - | + | - | - |
| | 22 | - | - | + | + | + | + | + | - | X | X | X |
| | 93 | - | - | - | / | - | X | X | X | X | X | X |
| | 26 | - | X | X | X | X | X | X | X | X | X | X |
| Vaginal | 89 | X | X | X | X | X | X | X | X | X | X | X |
| | 28 | X | X | X | X | + | + | X | X | X | X | X |
| | 90 | X | X | X | X | + | + | + | + | + | + | - |
| | 22 | X | X | X | X | + | + | + | + | X | X | X |
| | 93 | X | X | X | X | + | X | X | X | X | X | X |
| | 26 | X | X | X | X | X | X | X | X | X | X | X |

Symbols*: (x) = Not collected; (-) = Not detected; (/) = Suspect-positive; (+) = Positive, 38>Ct>31; (++) = Strong positive, Ct<31*

**Supplementary Table 6: Clinical monitoring criteria and scoring.**

| Parameter | Monitoring Criteria | Clinical Score Criteria | | | |
|---|---|---|---|---|---|
| | | **0** | **1** | **2** | **3** |
| Activity/Attitude | Depression, decreased alertness and/or responsiveness | Normal | Decreased alertness | Depression, Head down, unwilling to stand | Not responsive |
| Rectal temperature | Temperature above 104.5 F will be considered as fever | 100.4 - 102.9°F | 103.0-104.4°F | >104.5°F for 2+ days | >107.0°F |
| Appetite | Intake of food by an animal. | Normal | Reduced | Low | Zero (refuse feed and enrichment snacks) |
| Respiratory Signs | Sneezing, coughing, labored breathing, nasal discharge | Normal | Mild-Heavy breathing | Consistent heavy breathing, wide stance, abdomen expanding | Same as above, plus, wheezing |
| Mastitis | Abnormalities in the udder, such as swelling, heat, hardness, or pain | Normal | Mild swelling; Warm to touch; Firm; Sensitive | Swelling is significant; Red appearance and warm; Hard; Painful | Bulging, red/hot; Hard; Severe pain |
| Neurological | Observations on cow posture and gait/locomotion | Normal; Walking evenly | Walks unevenly | Lame (specific limb?) | Very lame; unresponsive to touch |
| Conjunctivitis | Observations on eye condition | Normal | Red eye; minor discharge | Red eye; moderate bi-lateral discharge | Red eye; Unable to open eye; Heavy ocular discharge |
